# Exploring the mechanisms of early acquired resistance to doxorubicin in melanoma spheroids

**DOI:** 10.1101/2025.04.07.646990

**Authors:** Giorgiana-Gabriela Negrea, Ilie Ovidiu Pavel, Loredana Balacescu, Bogdan-Razvan Dume, Emilia Licarete, Valentin-Florian Rauca, Laura Patras, Szilvia Meszaros, Stefan Dragan, Vlad Alexandru Toma, Manuela Banciu, Alina Sesarman

## Abstract

This study investigated the mechanisms underlying early settlement of doxorubicin (DOX) resistance in B16.F10 murine melanoma spheroids, following repeated exposure to a subinhibitory concentration of the drug. Melanoma spheroids were twice treated with DOX for 48h with a 48h recovery period, and changes in viability, growth, gene/protein expression, and enzyme activity were assessed using RNA-seq, RT-qPCR, western blot, protein array, and gelatinase assays. DOX exposure triggered a biphasic response in melanoma spheroids, with the initial exposure downregulating transcripts involved in cell cycle, DNA damage and repair responses, and suppressing of *TNF-α via NF-κB* and *mTORC1* stress response-related signaling pathways, indicating cell cycle arrest, enhanced DNA damage, and apoptosis resistance. Concurrently, upregulation of *Notch1*, and of angiogenic, adhesion, and ECM remodeling genes and proteins indicated early DOX-adaptive responses aimed at evading checkpoint arrest and increasing cell aggressiveness. A second DOX exposure amplified these responses in melanoma spheroids, leading to upregulation of some genes involved in cell cycle progression, DNA repair damage responses, along with increased *Aqp1*, *VEGF*, *Ackr3*, *MMP-2* expression, as well as elevated MMP-9 activity. Our results offer valuable insights into the molecular drivers of chemoresistance, revealing that early DOX-resistance in melanoma arises from adaptive mechanisms that support cell survival through enhanced angiogenesis and cell migration and motility capacity.

## Introduction

Melanoma tumors are very aggressive, with significant intrinsic heterogeneity, that favors the acquisition of resistance to various therapeutics and rapid metastatic progression ^1^. BRAF and MEK inhibitors are effective in BRAF-mutant melanoma, while immune checkpoint inhibitors have reshaped the treatment protocols for this disease, achieving long-term remission in a subset of patients ^2–5^. Doxorubicin (DOX), an anthracycline with limited use in frontline melanoma care, could be relevant, particularly for those patients who are not eligible or do not respond to the above-mentioned therapeutic approaches ^6,7^. Previous studies, including our own, demonstrated limited efficacy of DOX in melanoma models due to chemoresistance ^8–10^. Moreover, the low response rates and limited survival benefit ensured by liposomal DOX in metastatic melanoma ^11^ highlights the need for new therapeutic strategies to overcome drug resistance. DOX-based combination therapies, including ours ^12,13^, showed improved efficacy against melanoma in experimental studies, while others enhanced immunogenicity, sensitizing tumors to checkpoint inhibitors ^14^. Resistance to DOX can develop within weeks, driven mainly by ABCB5-mediated drug efflux and activation of survival signaling pathways ^15–18^. Exposure to DOX at sub-inhibitory concentrations (<IC50) induces transient drug-tolerant states through chromatin remodeling, stress responses, and dedifferentiation ^19^. creating favorable conditions that enable the selection of resistant cell subclones ^20,21^. In this study we explored the early mechanisms of chemoresistance in B16.F10 melanoma spheroids after repeated administration of a sublethal (IC_30_) concentration of DOX. Our results highlighted a two-phase adaptive response of B16.F10 melanoma spheroids to DOX. Particularly, mRNA/protein analyses evidenced an initial arrest of melanoma cells likely at G2/M checkpoint supported by downregulation of cell cycle, DNA damage and repair, and apoptotic genes, followed by survival-driven reprogramming, marked by upregulation of *Notch1* and of angiogenic, adhesion, and ECM remodeling genes and proteins. Re-exposure to DOX intensified some of these responses, increasing the expression of *Aqp1, VEGF, Ackr3*, *MMP-2* transcripts, and enhancing MMP-9 activity. Collectively, these findings revealed that melanoma cells withstand DOX-induced stress ^22^ by activating a coordinated mechanism that favor survival over cell death ^9,22,23^. Thus, combining DOX with selective inhibitors of Notch1, cIAP2, and/or with anti-metastatic drugs, might help overcome early resistance and suppress tumor progression^24–27^.

## Materials and Methods

### Murine melanoma spheroids formation

For spheroid formation ^28^, B16.F10 melanoma cells (5×10³/well; ATCC CRL-6457, Lonza Group, Switzerland) were seeded in ultra-low attachment 96-well plate, in 1.5% extracellular matrix (Sigma-Aldrich, Germany), using DMEM medium with supplements ^29,30^. Plates were centrifuged (15 min, 350×g) and incubated at 37°C with 5% CO₂. Spheroids images were taken daily, for diameter measurement, using a Zeiss Axio Vert.A1 microscope and SpheroidSizer (https://bit.ly/4hQBJZf).

### Drug treatment

Melanoma spheroids (day 4 post-seeding) were treated with DOX (Sigma-Aldrich, Germany) (0.156–10 µM), while untreated spheroids served as controls (CTR1,2). After 48h, cell viability was measured to determine IC_50_/IC_30_ values (**Figure 1A**) ^31^. For early resistance analysis, spheroids received two DOX (IC₃₀) exposures, separated by a 48h recovery phase (**Figure 1B**) ^32–34,33^. Spheroids were collected 48h after 1^st^ DOX exposure (DOX1, CTR1) for RNA/ protein extraction and remaining spheroids were re-exposed to DOX (DOX2) or left untreated (CTR2). Growth rates (k_DOX1,2_ and k_CTR1,2_) and resistance growth marker (k_DOX/_k_CTR_ ≥1=resistance to drug; <1=drug sensitivity) were calculated (**Figure 2A,B**) ^34^.

**Figure 1.**
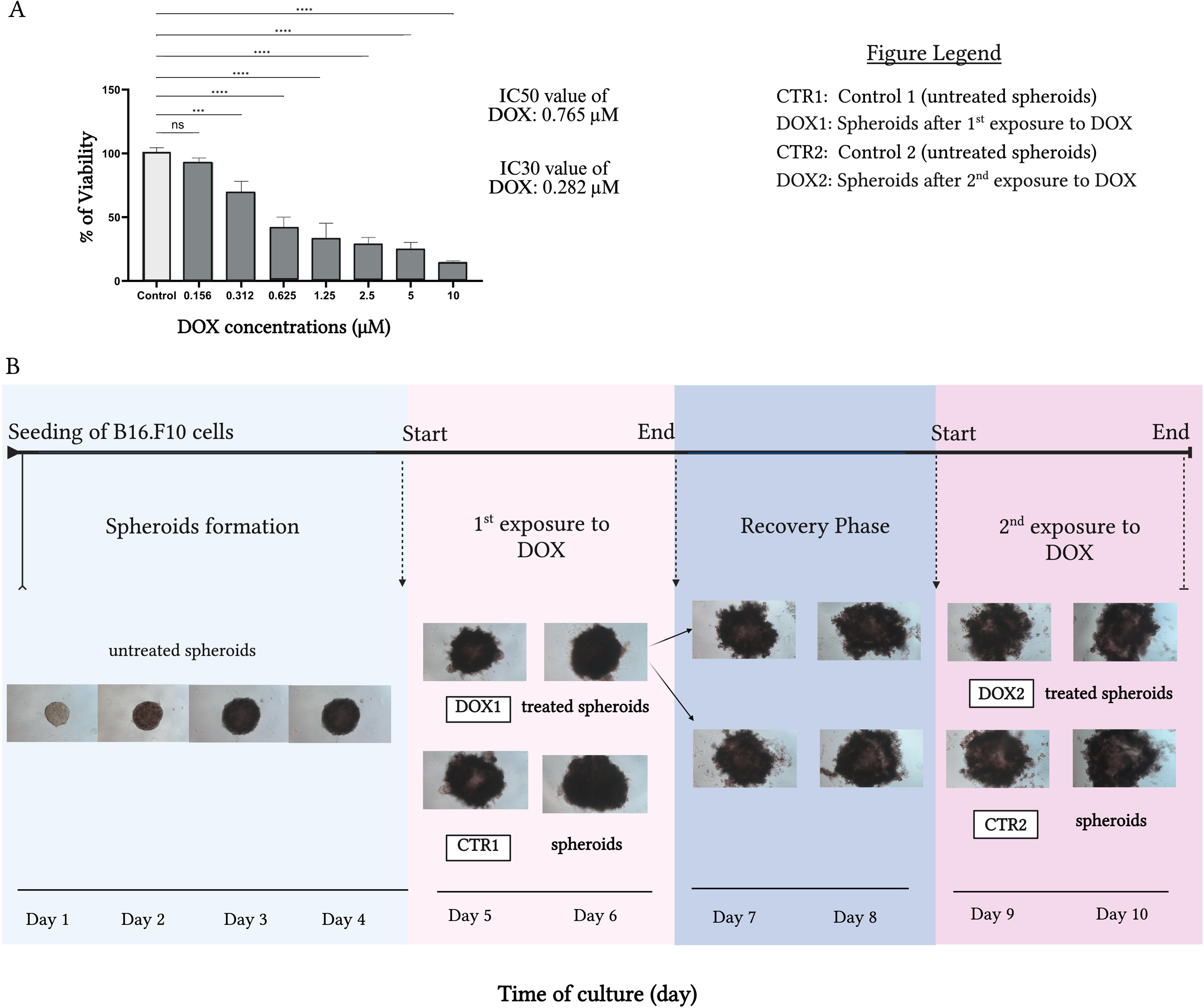
Cell viability, experimental timeline, and morphology of B16.F10 spheroids exposed to DOX. **A. Cell** viability (%) following DOX (conc. 0.156–10 µM) exposure measured by APH assay. **B.** Experimental timeline and representative spheroid images (5x objective, 50× magnification).

**Figure 2.**
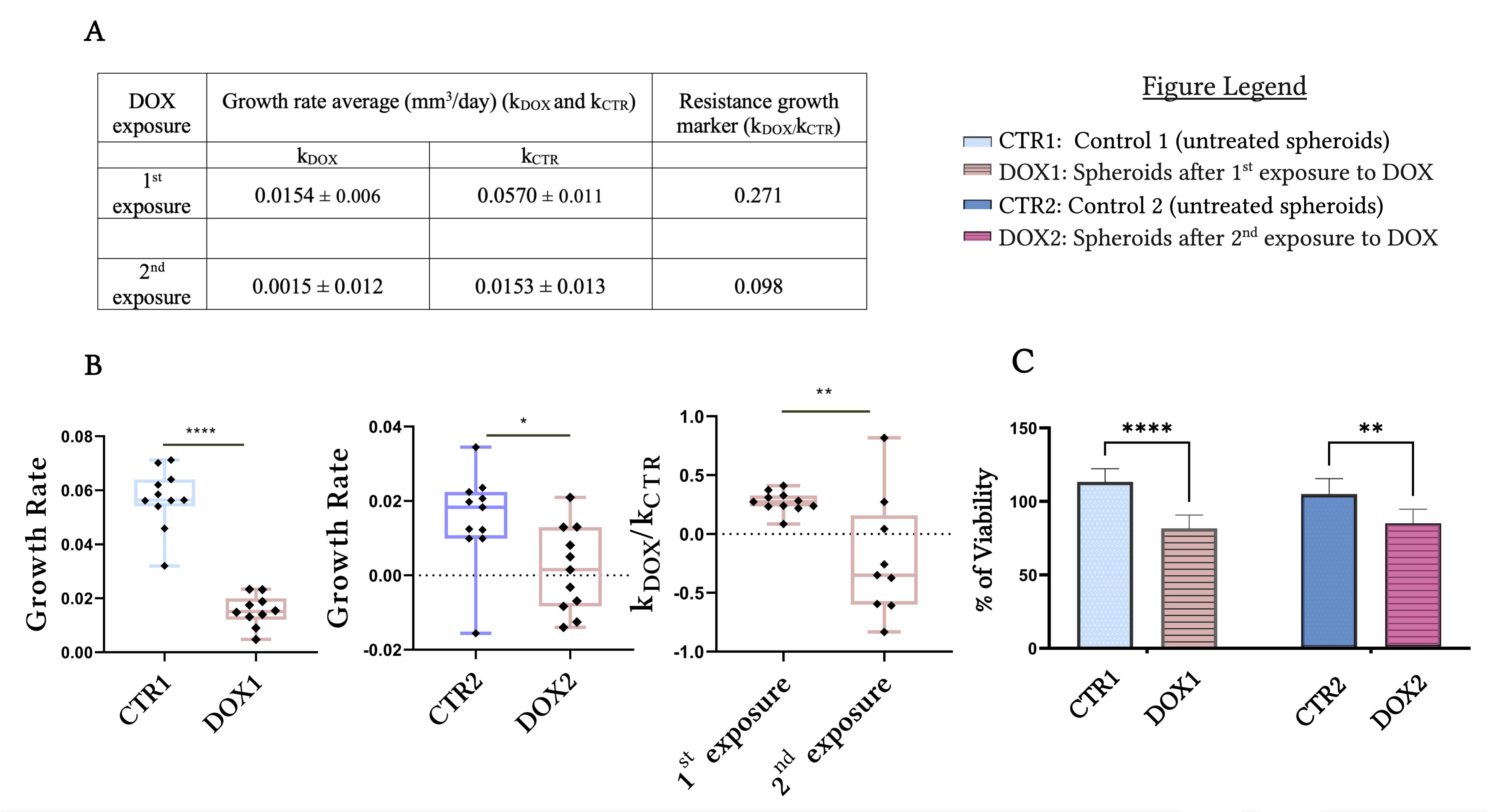
Growth rate and viability of B16.F10 spheroids treated with DOX. **A.** Growth rates during DOX exposures (K_DOX1,2_) versus controls (K_CTR1,2_), and resistance growth marker (K_DOX_/K_CTR_). **B.** Graphical representations of growth rates. **C.** Spheroids viability (%) after the 1^st^ and 2^nd^ DOX exposure (IC30), versus controls. Results are presented as mean ± SD of three independent experiments (n=9).

### Measurement of cell viability within the spheroids

To assess cell viability within the spheroids exposed to DOX, acid phosphatase (APH) assay was performed (Fluostar Omega, BMG-Labtech, Germany) ^35^. Cell viability in DOX-treated spheroids was expressed as a percentage relative to control spheroids. The experiments were conducted in triplicate. All data were expressed as the mean ± SD (n=9).

### Total RNA isolation

Total RNA was extracted from spheroids using the RNeasy Mini Kit (Qiagen, Germany) following the manufacturer’s protocol ^36^. RNA quality and integrity were assessed with a NanoDrop 1000 (Thermo Scientific, USA). Strand-specific cDNA libraries were prepared using the TruSeq Stranded mRNA Library Prep Kit (Illumina), followed by quality control (performed by Macrogen, Inc (www.macrogen-europe.com)).

### RNA-sequencing and data analysis

Libraries were sequenced on Illumina Paired-End-101 by Macrogen with two biological replicates per group ^37^. Reads were trimmed, aligned to the mouse genome using HISAT2 (v2.0.5), and transcripts assembled with StringTie-eB (v1.3.1). Transcript abundance was reported as raw counts and normalized (FPKM, CPM, TPM) ^38^. Low-quality transcripts were filtered, and data normalized with TMM in edgeR. Differential expression was assessed by exactTest with Benjamini-Hochberg correction ^39,40^. DEGs between DOX1 and CTR1 were defined by |FR| ≥ 2, FDR<20%, and p<0.05. Functional enrichment used g:Profiler (GO) and DAVID (KEGG), visualized in R (ggplot2) ^41^. Volcano plots and hierarchical clustering of Z-scores were generated using heatmap.2 in R. Z-scores were calculated as: z=(x-μ)/σ (*x*=gene count in a replicate, *μ*=mean, *σ*=standard deviation). Gene Set Enrichment Analysis GSEA with HALLMARK sets identified enriched pathways; significance: FDR<20%, p<0.05 ^42^. All analyses after sequencing were performed by the authors.

### Quantitative Real-Time PCR (RT-qPCR)

RT-qPCR was performed to validate RNA-seq results. Primer sequences are available in **Supplementary Table 1,** Repository ^43^ while full protocol details are available in the **Supplementary Material.**

### Spheroids lysates preparation

B16.F10 spheroids were washed, lysed in 250 μl RIPA buffer (Invitrogen) with proteases and phosphatases inhibitors. Protein content was measured by Bradford assay (Sigma-Aldrich, Germany) ^44^.

### Apoptosis array

Apoptotic protein expression was analyzed using 250 µg lysate and a 38-target antibody array (https://www.raybiotech.com/mouse-apoptosis-array-c1-aam-apo-1), following manufacturer’s instructions^45^. Immune complexes were detected by chemiluminescence (ChemiDoc, Bio-Rad, USA) and Image Lab 4.1 (Bio-Rad). Each sample was assessed in duplicate.

### Western blot analysis

pNF-κB expression was detected from 50 μg of protein per group via western blot, following a published protocol ^46^. Immune complexes were detected by chemiluminescence using ChemiDoc Imaging System (Bio-Rad, USA) and Image Lab Software 4.1 (Bio-Rad). Data from three independent experiments are presented as mean ± SD.

### Angiogenesis array

Angiogenic/inflammatory proteins were analyzed using 250 µg lysate, and a 24-target antibody array (https://www.raybiotech.com/mouse-angiogenesis-array-c1-aam-ang-1), following manufacturer’s instructions^9^. Immune complexes were detected by chemiluminescence (ChemiDoc, Bio-Rad, USA) and Image Lab 4.1 (Bio-Rad, USA). Each sample was assessed in duplicate.

### Gelatin zymography

MMP activity was measured from spheroid supernatants ^47^. 50 µg of protein per group were run on 7.5% gelatin-containing SDS-PAGE gels under non-reducing conditions ^9^. Enzyme activity was quantified by densitometry using Bio-Rad’s ChemiDoc (USA) and Image Lab 4.1. Experiments were performed in duplicate.

### Statistical analysis

Data are reported as mean ± SD. Statistical analyses were performed using GraphPad Prism 7. DOX IC50 values were calculated *via* nonlinear regression of dose-response curves; IC30 was calculated in Excel using: ICx=[(x/100-x)^1/Hillslope]*IC_50_ ^31^. Unpaired Student’s t-test was used to compare mRNA and protein expression in spheroid groups; two-way ANOVA was used for comparing the data obtained in protein arrays. Significance was set at p < 0.05 (*), with **p < 0.01, ***p < 0.001, ****p < 0.0001; ns = not significant.

## Results

### The effect of DOX on the viability and growth rate of B16.F10 melanoma spheroids

As shown in **Figure 1A** the viability of melanoma cells within the spheroids significantly decreased at concentrations >0.156 µM (p<0.001). The IC₅₀ and subinhibitory IC₃₀ values were calculated as 0.765 µM and 0.282 µM, respectively. Four-day-old melanoma spheroids were subjected to two consecutive exposures to DOX (IC₃₀), each followed by a 48h recovery phase, as illustrated in **Figure 1B** ^48^. Representative images (**Figure 1B** and **Supplementary Material-Figure 1**) showed rounded shape spheroids of 400-700 µm diameters over the 10-day observation period. DOX significantly reduced spheroid growth rate (DOX1 vs. CTR1, p<0.0001; DOX2 vs. CTR2, p<0.01, **Figure 2A,B**), likely due to reduced B16.F10 cell viability (**Figure 2C**). A resistance marker below 1 (k_DOX2/_k_CTR2_=0.098) confirmed sensitivity to repeated DOX exposure (**Figure 2A,B**) ^34^.

### Transcriptional changes associated with the early response of melanoma spheroids to DOX

To explore early molecular changes associated with drug resistance in B16.F10 melanoma spheroids after a single DOX exposure, RNA-seq analysis was performed. Using FR ±2, FDR <20%, and p <0.05, a total of 1040 DEGs were identified (520 upregulated, 520 downregulated; **Supplementary Table 2,** Repository ^43^). Results were visualized *via* unsupervised hierarchical heatmap (**Figure 3A**), volcano plot (**Figure 3B**), and DEGs distribution bar plot (**Figure 3C**). GO enrichment analysis (**Figure 3D**) revealed suppressed processes including cell cycle, DNA damage response, DNA repair, and angiogenesis (**Supplementary Table 3.1**, Repository ^43^). KEGG pathway analysis (**Figure 3E** showed seven downregulated pathways (e.g., cell cycle, DNA replication, TNF signaling) and two upregulated (p53 and NOD-like receptor signaling) (**Supplementary Table 3.2**, Repository^43^). GSEA (**Figure 4A**) identified six enriched gene sets, with five negatively enriched (e.g., G2M checkpoint, E2F targets, TNFA *via* NF-κB) and one positively enriched (Interferon-α response), with expression profiles shown in **Figure 4B-G; Supplementary Material-Figure 2; Supplementary Table 4**, Repository ^43^).

**Figure 3.**
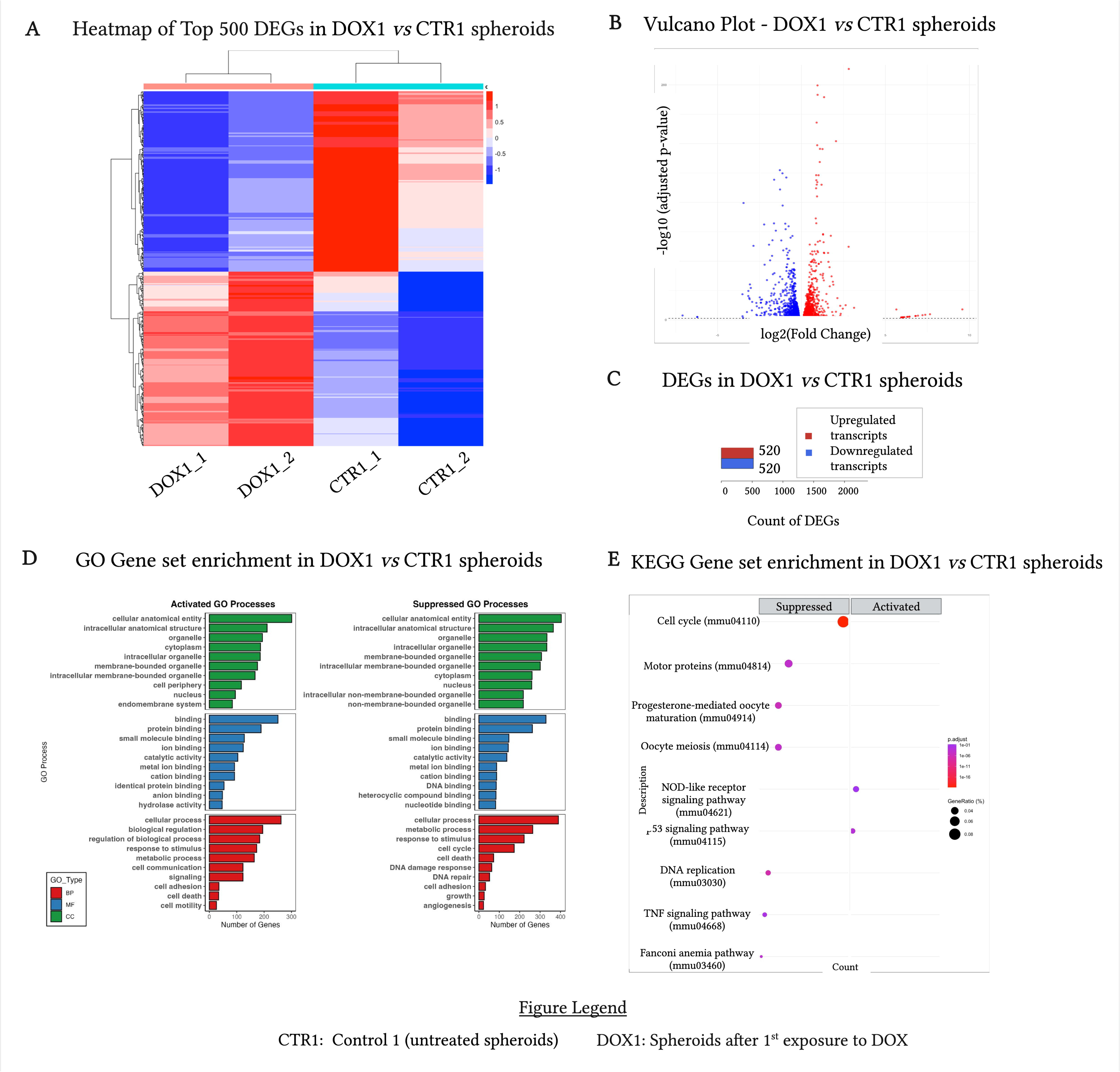
RNA-seq analysis of spheroids after 1st DOX exposure vs. control. **A.** Heatmap of top 500 DEGs (unsupervised clustering) based on normalized FPKM; red=upregulated, blue=downregulated (DOX1 vs. CTR1). **B.** Volcano plot. **C.** DEGs bar plot. **D.** GO enrichment: activated (left) vs. suppressed (right); BP (red), CC (green), MF (blue). **E.** KEGG pathways: activated (right) vs. suppressed (left).

**Figure 4.**
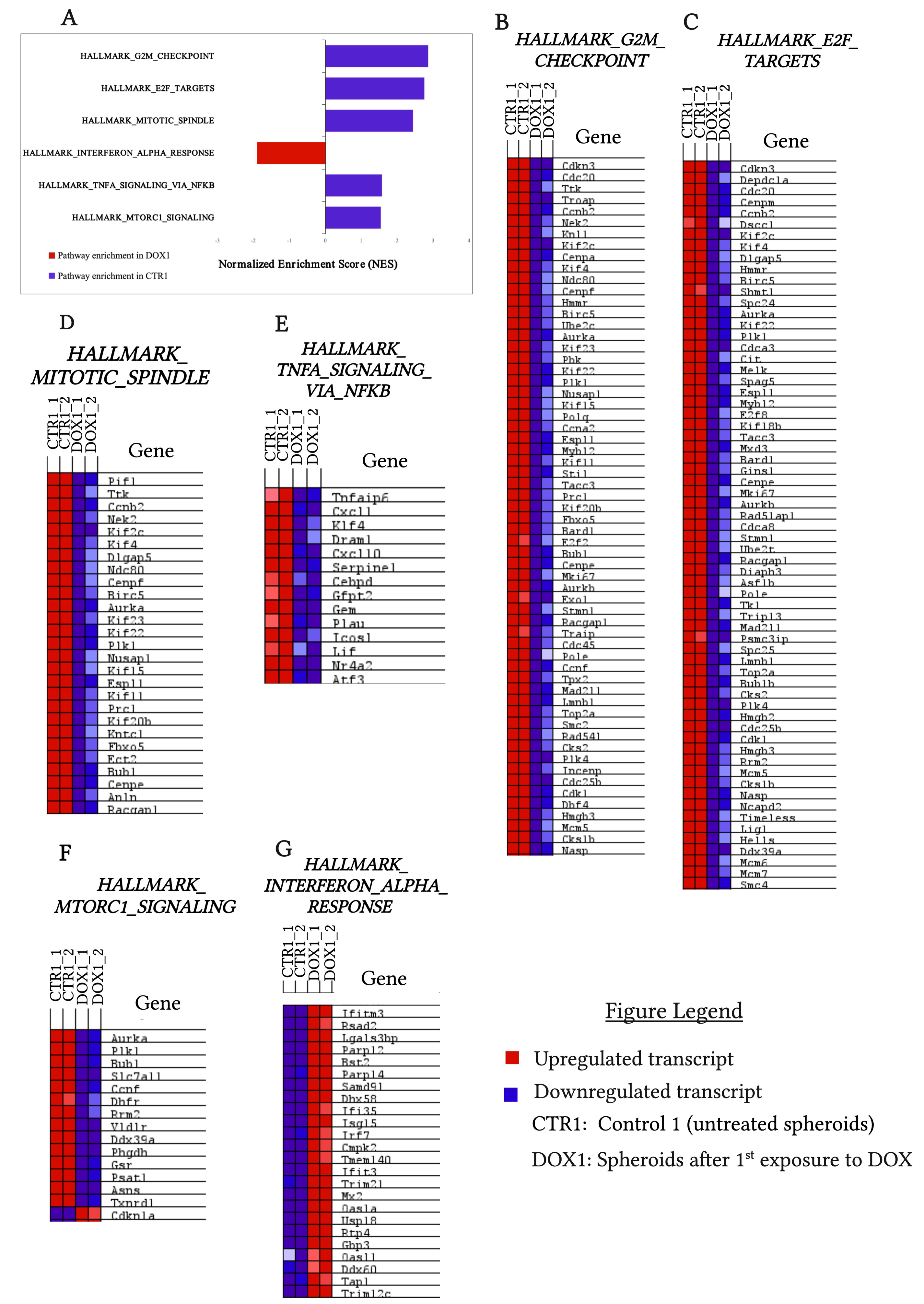
GSEA Hallmark analysis of spheroids after 1^st^ DOX exposure vs. control Gene sets with FDR<20% and p<0.05 are shown. **A.** Bar plot of enriched pathways: positive Normalized Enrichment Score (NES)=enrichment in CTR1; negative NES=enrichment in DOX1. **B-G.** Heatmaps of selected gene sets comparing CTR1 (left) and DOX1 (right). Expression levels: red (high) to dark blue (low).

#### Repeated DOX exposure triggered initial cell cycle arrest followed by partial reactivation of cell cycle progression in melanoma spheroids

To explore early transcriptional alterations in response to 1^st^ DOX exposure, hierarchical clustering of DEGs was performed, revealing significant alteration is key processes, cell cycle, DNA damage/repair, apoptosis, angiogenesis, and cell migration/motility capacity in DOX-exposed spheroids (DOX1) compared to control (CTR1) (**Figure 5A**; **Supplementary Table 5**, Repository^43^). Cell cycle pathways exhibited the most pronounced changes with 189 DEGs (**Figure 5B-D**; GSEA results in **Figure 4A-D**; **Supplementary Material-Figure 2; Supplementary Table 5**, Repository ^43^). Notably, RNA-seq revealed downregulation of core mitotic regulators such as *CDK1 (p<0.001), Nek2 (p<0.001), Bub1 (p<0.001), Mad2l1 (p<0.001), and Ticrr (p<0.001), while CDKN1A/p21 (p<0.001)* and *Psrc1 (p<0.001)* were upregulated (**Supplementary Table 5**, Repository ^43^). RT-qPCR confirmed the RNA-seq results, showing consistent transcriptional repression of some genes that were selected for validation such as *CDK1* (p<0.05)*, Nek2* (p<0.05)*, CDC20* (p<0.01)*, Bub1* (p<0.05)*, Mad2l1* (p<0.05), and *Ticrr* (p<0.05) after the 1^st^ DOX exposure (**Supplementary Material-Figure 3A-G**) likely contributing to DOX initial cytotoxic effects (**Figure 2C**). Following the 2^nd^ exposure, RT-qPCR showed that some of these genes remained suppressed (*CDC20*, p<0.01), few showed moderate upregulation (*Bub1,* p<0.05), while others had expression levels approaching those observed in controls (CTR1 and CTR2) (*CDK1,* p>0.05*; Nek2,* p>0.05*; Mad2l1*, p>0.05) (**Supplementary Material-Figure 3A-G**). Collectively, our findings suggested that repeated exposure to DOX might have supported early adaptive transcriptional changes that re-initiated cell cycle progression.

**Figure 5.**
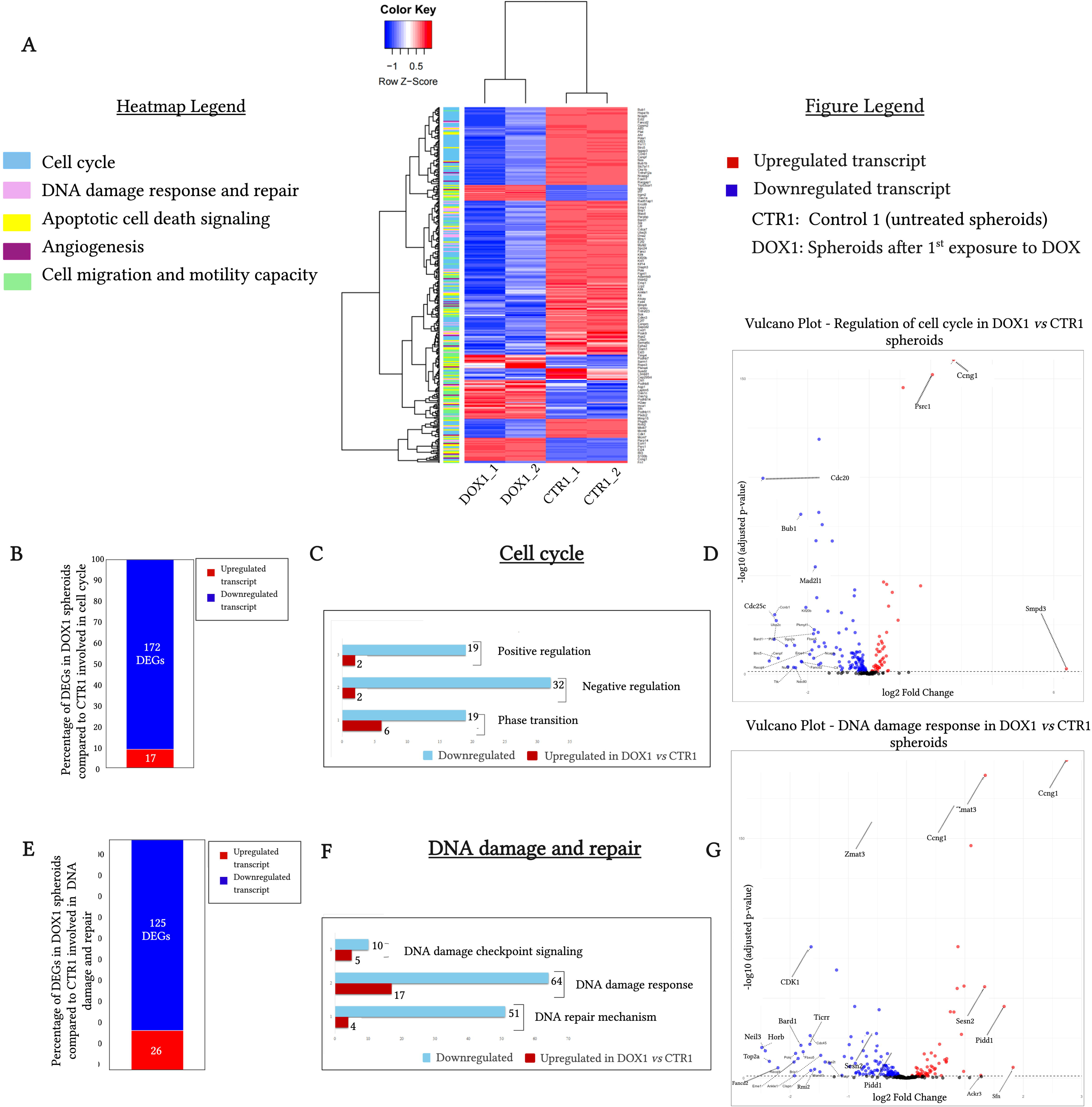
RNA-seq analysis of DEGs related to cell cycle and DNA damage and repair processes in DOX1 vs. CTR1 spheroids. **A.** Heatmap (unsupervised clustering) based on normalized FPKM; red=upregulated, blue=downregulated (DOX1 vs. CTR1). **B-D.** Cell cycle-related transcripts. **E-G.** DNA damage/repair related transcripts. Color code: Red=upregulated; Blue=downregulated in DOX1 vs. CTR1. Generated using R and Excel.

#### Initial DOX exposure altered DNA repair and damage transcripts in melanoma spheroids, while a second one initiated genome stability recovery

RNA-seq revealed that a single exposure to DOX significantly impaired DNA repair and damage response mechanisms in melanoma spheroids (**Figure 5E-G; Supplementary Table 5,** Repository ^43^). A total of 150 DNA damage–related DEGs were altered, with prominent downregulation of *Top2A* (p<0.0001) or *Rmi2* (p<0.001). 64 transcripts involved in DNA-damage sensing pathways, including *Cdc45 (p<0.0001), Foxm1, Fancd2, Exo1, Brip1, Bard1, and Rad51* (all p<0.001) were downregulated and 17 transcripts such as *Trex1, EI24, and Sesn2* upregulated (all p<0.001). Additionally, 51 DEGs linked to DNA-repair showed reduced expression, including *Polq, Parpbp,* and *Fanci* (p<0.001), while 4 were upregulated. RT-qPCR confirmed RNA-seq findings, with significant downregulation of *Rmi2* and *Histone 1X* (both p<0.05) after the 1^st^ DOX exposure. Interestingly, a 2^nd^ DOX exposure induced an increase in *Rmi2 expression* (p<0.05), while *Histone 1X* levels remained unchanged, resembling control (**Supplementary Material-Figure 3H,I**), indicating that while DOX initially suppressed DNA repair gene expression, repeated exposure might have promoted gene-specific transcriptional adaptations supporting genomic stability.

#### Repeated DOX administration failed to activate pro-apoptotic transcriptional programs in melanoma spheroids

Apoptosis-related pathways showed significant transcriptional changes with 106 DEGs identified (**Figure 6A-C**). Both intrinsic and extrinsic apoptosis signaling were suppressed in spheroids after a single DOX exposure, indicated by downregulation of *survivin* (*Birc5*, p<0.001) and of *BOK* (p<0.001), alongside upregulation of *PIDD1* (p<0.001) (**Supplementary Table 5**, Repository^43^). Moreover, negative enrichment of TNFA_SIGNALING_VIA_NFKB (**Figure 4E**) suggested suppressed transcription of NF-κB-mediated anti-apoptotic regulators. RT-qPCR confirmed significant upregulation of *stratifin* (*Sfn*; p<0.05) after initial DOX exposure, while *Bax*, *Bid*, and *Bcl-2* expression remained unchanged (p>0.05) compared to control. After the 2^nd^ DOX exposure, *Bcl-2* showed a slight expression reduction (p<0.05), whereas *Sfn*, *Bax*, and *Bid* remained unchanged compared to control (p>0.05) (**Supplementary Material-Figure 3J-M**). Overall, these changes in apoptotic transcripts indicated limited apoptosis following both DOX exposures, potentially conferring melanoma spheroids a survival advantage (**Figure 6A-C; Supplementary Material-Figure 3J-M; Supplementary Table 5,** Repository ^43^).

**Figure 6.**
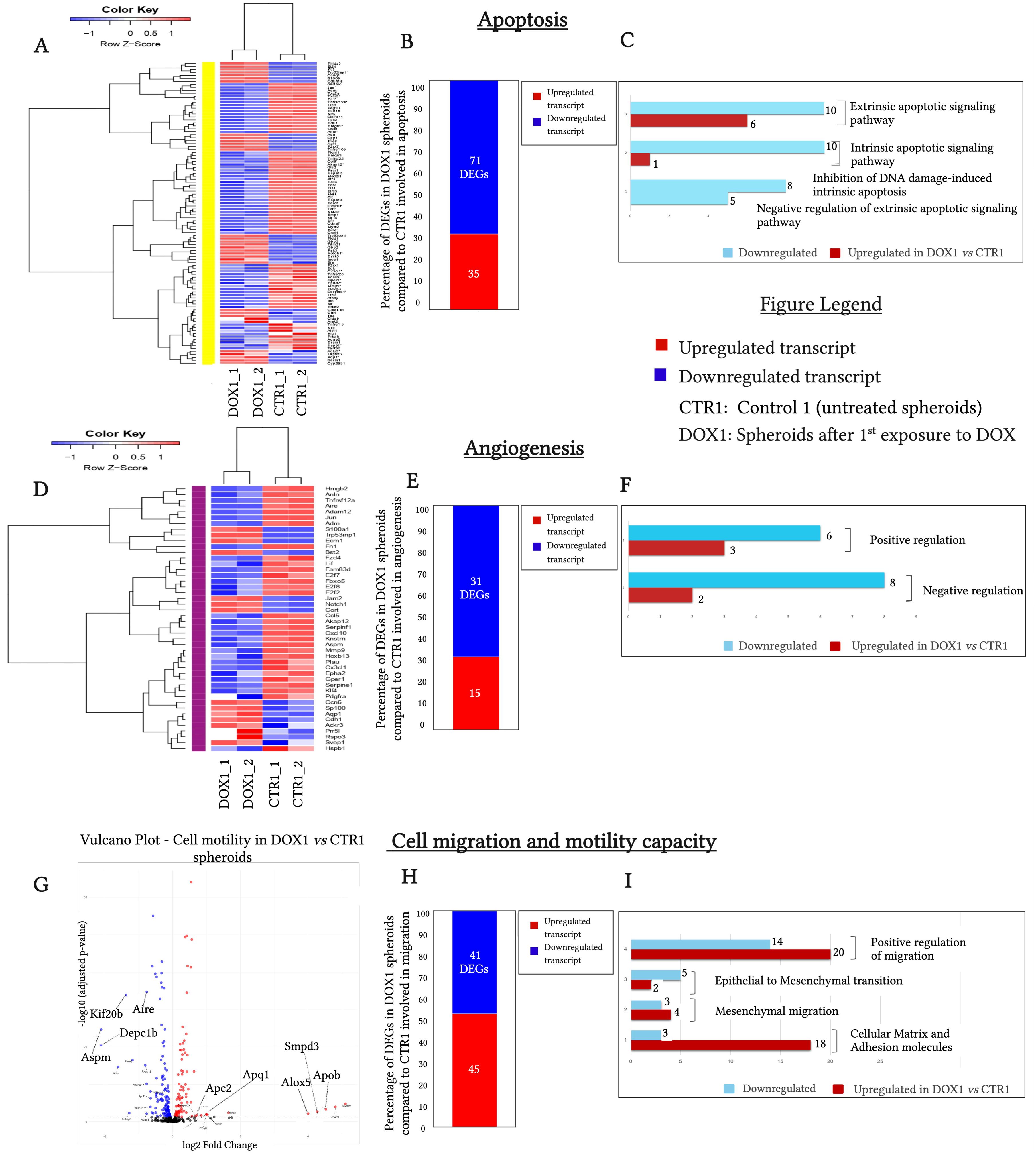
RNA-seq of DEGs related to apoptosis, angiogenesis, and migration in spheroids after 1st DOX exposure vs. control. **A-C.** Apoptosis-related transcripts. **D-F**. Angiogenesis-related transcripts. **G-I.** Cell migration and motility capacity related transcripts. Heatmaps (unsupervised clustering) based on normalized FPKM; red=upregulated, blue=downregulated (DOX1 vs. CTR1). Generated using R and Excel.

#### Biphasic modulation of angiogenesis-related transcripts in melanoma spheroids, following repeated DOX exposure

Although GSEA showed negative enrichment of TNFA_SIGNALING_VIA_NFKB and MTORC1_SIGNALING angiogenic/inflammatory pathways (**Figure 4E,F**) after a single exposure of spheroids to DOX, RNA-seq revealed a total of 46 angiogenesis-related DEGs (31 downregulated, 15 upregulated) (**Figures 6D-F; Supplementary Table 5**, Repository ^43^). These included upregulation of pro-angiogenic *Bst2* and *Svep1* (p<0.01), and downregulation of vascular stabilizers *Epha2* and *Ccn6* (p<0.01). RT-qPCR showed a slight VEGF downregulation after initial DOX (p<0.05), followed by significant upregulation upon re-exposure of spheroids to DOX (p<0.05) (**Supplementary Material-Figure 3N**). *Aqp1*, involved in vascular permeability and cell migration, was consistently upregulated after both exposures of melanoma spheroids to DOX compared to control (p<0.05) (**Supplementary Material-Figure 3O**). Thus, these results are indicative of a two-phase angiogenic response triggered by DOX in melanoma spheroids, beginning with suppression and followed by activation of vascular remodeling and stress-adaptive transcriptional programs.

#### DOX at subinhibitory concentration favored migratory and motility gene expression in melanoma spheroids

Cell migration and motility-related genes (96 DEGs) were altered after a single exposure of spheroids to DOX (**Figure 6G-I; Supplementary Table 5**, Repository ^43^). Several protocadherins (*Pcdhβ5, β7, β8; p<0.001*) and adhesion-related transcripts were upregulated, while *Nectin4*, *Cdh6*, and *Itga3* were significantly downregulated (p<0.001) (**Figure 6G-I**). DOX also strongly suppressed the expression of *MMP-9* (2-fold, p<0.001), a key EMT regulator and upregulated that of *MMP-15* (p<0.001), of *Notch1* (p<0.001), and of other aggressiveness markers such as *Tnfrsf19, Plxdc2, IL-16, and Angptl2* (**Figure 6G-I, Supplementary Table 5**, Repository ^43^). RT-qPCR confirmed slight upregulation of *Ackr3* (p<0.05), a receptor linked to metastasis and downregulation of *MMP-9* (p<0.05), while *MMP-2* remained unchanged (p>0.05) in spheroids exposed once to DOX vs. control ^49^. Upon second DOX exposure, *Ackr3* and *MMP-2* were significantly upregulated (p<0.05), while *MMP-9* suppression was maintained (p<0.05), indicating a partial reactivation of transcription of pro-invasive genes (**Supplementary Material-Figure 3P-R**).

### Protein-level validation of transcriptomic changes induced by DOX in melanoma spheroids

To assess if DOX-induced mRNA changes (**Supplementary Table 5**, Repository ^43^) persist at the protein level, key proteins from the most affected pathways (apoptosis, angiogenesis and aggressiveness) were measured in B16.F10 spheroids (**Figure 7**).

**Figure 7.**
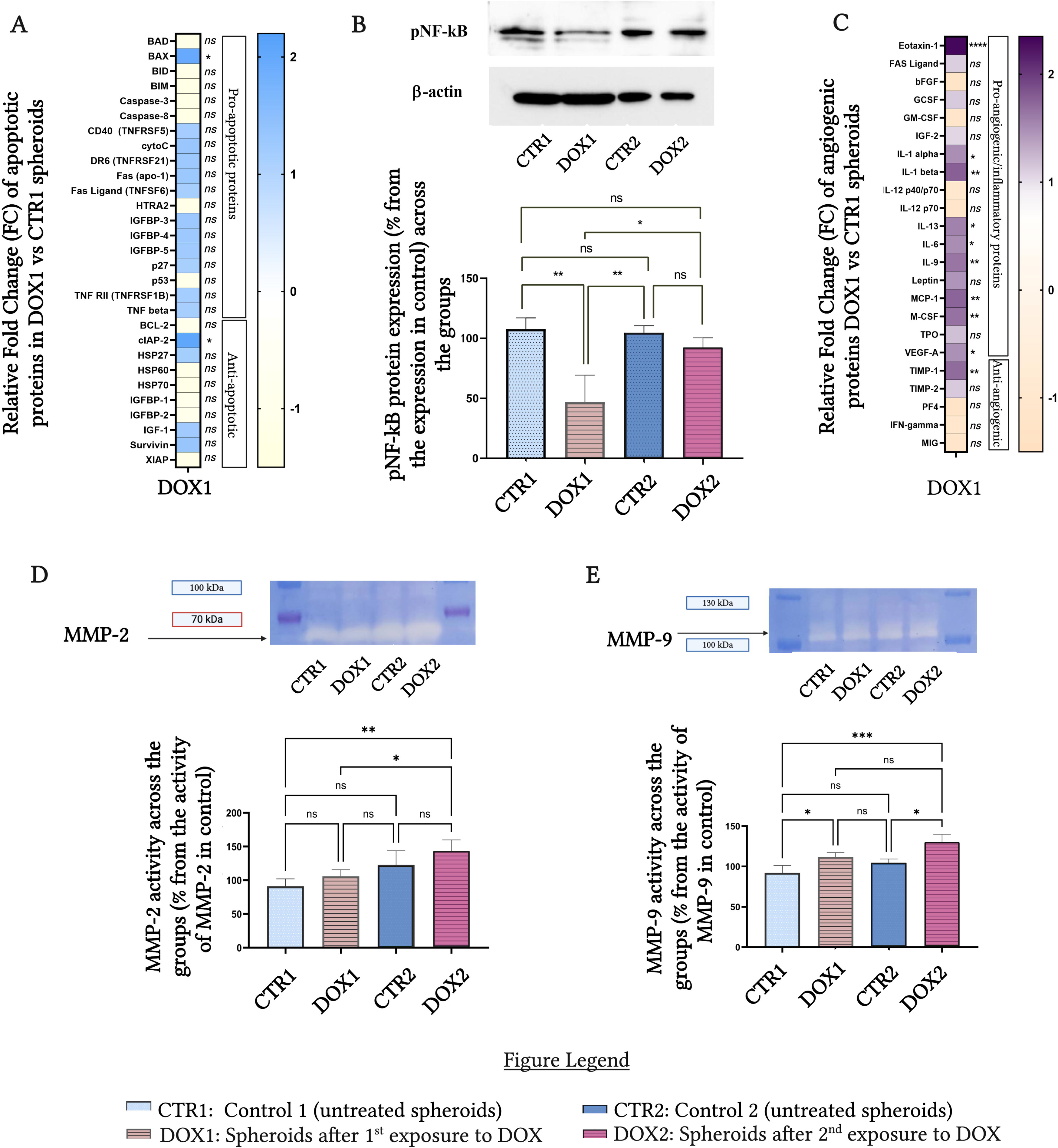
Effects of repeated DOX exposure on protein expression/enzyme activity in melanoma spheroids. **A.** Fold changes in apoptotic protein expression (DOX1 vs. CTR1): decreases (yellow, 0 to −1), increases (blue, 0 to +2); mean ± SD (n=2). **B.** Quantification of pNF-κB levels by Western blot after 1^st^ (DOX1) and 2^nd^ (DOX2) DOX exposure; β-actin as loading control; mean ± SD (n=3). **C**. Fold changes in angiogenic/inflammatory protein expression (DOX1 vs. CTR1): decreases (yellow), increases (purple); mean ± SD (n=2). **D, E.** Gelatin zymography of MMP-2 (**D**) and MMP-9 (**E**) activity after 1st and 2nd DOX exposure vs. controls; mean ± SD (n=2).

#### A single DOX exposure failed to induce apoptosis in melanoma spheroids

Our results demonstrated alteration of the expression level of several apoptotic proteins in DOX-exposed vs. to control spheroids (**Figure 7A**; **Table 1**). Among the 38 proteins tested, 9 proteins were not expressed in sufficient amount to be detected in spheroids lysates, namely BCL-W, IGFBP-6, IGF-2, p21, CD40L, TNF RI, TNF-α, TRAIL R2 and SMAC. Of the detected apoptosis-related proteins, only pro-apoptotic Bax and anti-apoptotic cIAP2 proteins were significantly upregulated (2-fold, p<0.05) in spheroids, after the first DOX exposure, while all the other proteins investigated showed no relevant changes (**Figure 7A**; **Table 1**) compared to their control level. These results, along with unchanged mRNA levels (**Supplementary Material-Figure 3K-M**) for most apoptotic proteins in DOX exposed spheroids, suggested that a single DOX exposure did not fully activate apoptosis in melanoma spheroids, likely due to adaptive survival mechanisms.

**Table 1.** Apoptosis protein array results. Level of expression of pro/anti-apoptotic proteins in single DOX exposured spheroids compared with the levels of the same proteins in control (untreated) spheroids. Results are presented as fold changes of reduction in protein levels ranging from 0 (white) to −1 (yellow), or as fold changes of increase in protein production ranging from 0 (white) to 2 (blue).

|  | Protein name | Fold change (FC) of inhibition (-) and stimulation (+) of proteins in spheroids after 1<sup>st</sup> exposure to DOX |
| --- | --- | --- |
| Pro-apoptotic proteins | BAD | 0.8719±0.2587 (ns) |
|  | BAX | 3.9098±0.1818 (*) |
|  | BID | 1.7546±0.2080 (ns) |
|  | BIM | 1.6089±0.0877 (ns) |
|  | Caspase-3 | 1.6421±0.3527 (ns) |
|  | Caspase-8 | 0.8086±0.0087 (ns) |
|  | CD40 (TNFRSF5) | 2.3207±1.1964 (ns) |
|  | cytoC | 2.6378±0.0353 (ns) |
|  | DR6 (TNFRSF21) | 2.5238±0.3125 (ns) |
|  | Fas (apo-1) | 2.5883±0.0290 (ns) |
|  | Fas Ligand (TNFSF6) | 2.2934±0.3596 (ns) |
|  | HTRA2 | 1.7063±0.1943 (ns) |
|  | IGFBP-3 | 1.2782±0.4907 (ns) |
|  | IGFBP-4 | 1.2649±0.5796 (ns) |
|  | IGFBP-5 | 1.2206±0.2061 (ns) |
|  | p27 | 1.1217±0.2385 (ns) |
|  | p53 | 0.9119±0.5996 (ns) |
|  | TNF RII (TNFSF1B) | 1.1745±2.3490 (ns) |
|  | TNF b | 1.4982±2.9964 (ns) |
| Anti-apoptotic proteins | BCL-2 | 1.9620±0.1855 (ns) |
|  | cIAP-2 | 4.1014±1.1240 (*) |
|  | HSP27 | 2.3136±0.2804 (ns) |
|  | HSP60 | 1.6628±0.4102 (ns) |

#### Enhanced expression of pro-angiogenic/pro-inflammatory protein in melanoma spheroids following a single DOX exposure

NF-κB is a key transcription factor involved in the survival of melanoma cells following chemo/immunotherapeutic stress including the modulation of angiogenesis and inflammation ^50^. Although GSEA showed suppression of the TNFA_SIGNALING_VIA_NFKB pathway (**Figure 4E**), except for NF-κB mRNA level, the expression levels of pNF-κB protein were reduced by 50% after initial exposure of spheroids to DOX (p<0.01) compared to CTR1, while returning to control (CTR2) levels after the second DOX administration to spheroids (p>0.05) (**Figure 7B**). Given RNA-seq evidence of modulated angiogenesis-related transcripts (**Figure 6D-F; Supplementary Table 5,** Repository ^43^), we further analyzed 24 angiogenic/inflammatory proteins *via* array, likely modulated by NF-kB ^51^. DOX increased (by 2-fold) the expression level of pro-angiogenic/inflammatory proteins-eotaxin-1, IL-1α, IL-1β, MCP-1, IL-6, IL-9, IL-13, M-CSF, VEGF-A (p<0.05–0.001), and of anti-angiogenic TIMP-1 (1.7-fold, p<0.01), while other proteins expression level remained unchanged (p>0.05)compared to control (**Figure 7C**, **Table 2**). Despite transcript-level suppression of angiogenic/anti-inflammatory genes (**Figure 6D-F**), these protein-level modification suggested that selective preservation of these key proteins through post-transcriptional regulation occurred in melanoma cells.

**Table 2.**
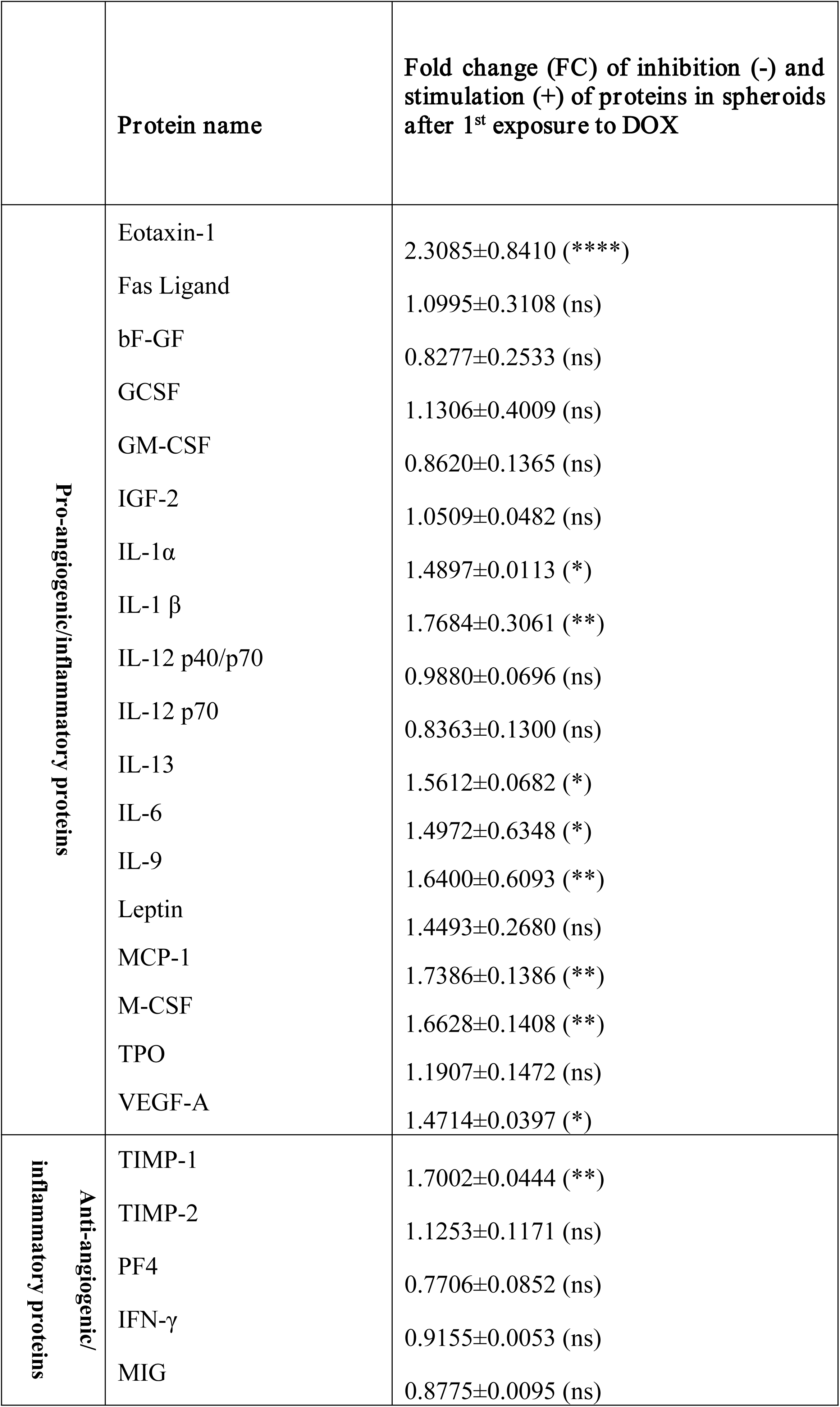
Angiogenesis protein array results. Level of expression of angiogenic/inflammatory proteins in spheroids after first exposure to DOX, compared with the levels of the same proteins in control (untreated) spheroids. Results are presented as fold changes of reduction in protein levels ranging from 0 (white) to −1 (yellow), or as fold changes of increase in protein production ranging from 0 (white) to 2 (purple).

#### DOX activated enzymes linked to tumor aggressiveness in melanoma spheroids

Since many NF-κB-regulated angiogenic and inflammatory proteins promote melanoma aggressiveness ^51^ and RNA-seq revealed altered expression level of cell migration/motility capacity-related transcripts in spheroids after a single DOX exposure (**Figure 6G-I; Supplementary Table 5**, Repository ^43^), we assessed the activity of the invasion markers, MMP-2 and MMP-9 ^52^. MMP-2 activity remained unchanged after 1^st^ DOX exposure (p>0.05; **Figure 7D**), while that of MMP-9 increased by 15% (p<0.05; **Figure 7E**) compared to control, indicating selective gelatinase activation. After the 2^nd^ DOX exposure, MMP-2 activity increased by 20% vs. DOX1 (p<0.05), but remained unchanged vs. CTR2 (p>0.05), and MMP-9 remained elevated when compared to it activity in CTR2 (p<0.05) with no difference between exposures (p>0.05). Together, the increased expression of pNF-κB in spheroids exposed twice compared to those exposed once to DOX and the upregulation of angiogenic and migration-related genes/proteins in spheroids receiving a single DOX administration (**Figures 4E; 5D–I; 7B,E; Supplementary Material-Figure 3R; Supplementary Table 5,** Repository ^43^) suggested a compensatory pro-survival response, likely modulated by NF-κB, that promoted angiogenesis, invasion, and chemoresistance ^51^.

## Discussion

Adaptive responses to chemotherapeutics are crucial for identifying early biomarkers of chemoresistance, enabling timely interventions, adjusting dosing strategies, and for the design of new combination therapies that prevent stable resistance and improve clinical outcomes ^53^. This study aimed to identify the mechanisms linked to the early settlement of resistance in B16.F10 murine melanoma spheroids, following repeated exposure to a subinhibitory concentration (IC_30_) of DOX. Thus, we noticed that an initial exposure of B16.F10 melanoma spheroids to DOX, led to downregulation of *Top2A* mRNA without affecting *ABC* transporters genes involved in DOX-efflux (**Supplementary Table 5,** Repository ^43^)^54^. This likely disrupted cell cycle dynamics, as pathways linked to proliferation and survival, identified by RNA-seq and GSEA analyses, were suppressed **(Figures 4A-F and 5A-D; Supplementary Material-Figure 3A-F; Supplementary Table 5,** Repository ^43^). This response was accompanied by a significant reduction in DNA repair capacity, particularly in homologous recombination (HR) and topoisomerase-induced damage mRNA **(Figure 5E-G; Supplementary Material-Figure 3G-I; Supplementary Table 5,** Repository ^43^) ^55,56^, likely suggesting that DOX exposure triggered cell cycle checkpoint arrest ^57^. As HR occurs during the S and G2 phases, melanoma cells, unable to enter mitosis, likely accumulated at G2/M checkpoint (**Supplementary Material-Figure 3A-F; Supplementary Table 5,** Repository ^43^)^58^. This is supported by downregulation of *Foxm1*, *Fanci*, and *DNA polymerases mRNA*, suggesting impaired DNA repair and reduced cell viability after DOX exposure (**Figure 2C**; **Supplementary Table 5,** Repository ^43^)^59^. Previous studies have demonstrated that shortly after exposure to cytotoxic drugs, cancer cells re-entered the cell cycle, leading to tumor relapse ^60^. We confirmed partial reactivation of cell cycle regulatory, and of DNA-damage and repair pathways, respectively, in melanoma spheroids, following the second DOX exposure (**Supplementary Material-Figure 3A-I**) ^61^, as a part of an adaptive transcriptional response that likely enabled melanoma cells to bypass the cell-cycle checkpoints ^62^, while still preserving drug sensitivity towards DOX (**Figure 2**) ^63^. Under conditions of pronounced DNA-damage and impaired repair capacity, cells attempting mitosis are usually susceptible to genomic instability, and prone to cell death ^64^. Interestingly, we demonstrated that DOX failed to fully activate apoptosis at transcriptomic or protein level in melanoma spheroids exposed once to DOX (**Figures 6A-C, 7A; Supplementary Material-Figure 3J-M; Table 1**;), despite the cell line maintaining functional wild-type p53 ^65^. Moreover, inhibition of NF-κB downstream of TNF-α signaling would typically enhance apoptosis mediated by this pathway (**Figure 4E**), yet the two-fold increase in the expression level of Bax (p<0.05) protein may have been counteracted by enhanced expression of cIAP2 protein (p<0.05), suggesting incomplete apoptosis (**Figure 7A**, **Table 1**) ^66^.

Our findings support a scenario where DOX-exposed melanoma cells bypassed cell cycle arrest, possibly engaging error-prone DNA repair mechanisms, as indicated by *Rad51* downregulation (**Supplementary Table 5,** Repository ^43^) ^67,68^, while a second exposure to DOX, equally failed to boost apoptosis, likely due to simultaneous *Bcl-2* and *cyclin G1* downregulation (p<0.05) (**Supplementary Material-Figure 3G,M**). Collectively, these findings suggested the settlement of early DOX resistance in melanoma spheroids via cell-cycle adaptation and apoptotic escape, enabling cell survival despite severe DNA-damage and impaired repair mechanism (**Supplementary Material-Figure 3A-M**) ^69^. Notably, this adaptive response included a biphasic regulation of angiogenesis and invasiveness, where initial DOX exposure induced transcriptional suppression of pro-angiogenic/pro-invasive transcripts, while post-transcriptional mechanisms seemed to preserve their protein expression (**Figures 6D-I and 7C-E; Supplementary Table 5,** Repository ^43^). Interestingly, some of these pro-angiogenic/pro-invasive transcripts, were upregulated upon repeated exposure of spheroids to sublethal DOX concentration (**Supplementary Material-Figure 3N-P**). Moreover, RNA-seq and GSEA further emphasized that an initial exposure of melanoma spheroids to DOX, suppressed mTORC1 and TNF-α via NF-κB signaling alongside upregulation of *Notch1* mRNA (p<0.001), possibly indicating a shift of melanoma cells towards an invasive, Notch1-driven phenotype (**Figures 4A,E,F and 6D-I; Supplementary Material-Figure 3N-R; Supplementary Table 5,** Repository ^43^) ^68,70–72^. Moreover, the increased levels of pro-invasive markers, including of NF-κB and MMP-9 activity, after repeated exposure of melanoma spheroids to DOX (**Figure 7B-E**; **Table 2**), reinforced early chemoresistance settlement and transition to an invasive cell phenotype ^73,74^.

In conclusion, early chemoresistance in B16.F10 melanoma spheroids likely resulted from DOX-induced stress that impaired cell cycle progression and compromised DNA-damage and repair mechanisms, triggering adaptive responses that enabled apoptosis evasion, *Notch1* activation, angiogenesis, and ECM remodeling, leading to acquisition of a more invasive, resistant cell phenotype. Combination strategies involving DOX and Notch1 inhibitors (γ-secretase inhibitors), LCL161 (cIAP inhibitors), and/or Batimastat (anti-metastatic agent) may enhance apoptosis, reduce transient resistance and limit tumor aggressiveness ^24–27^. While this study provides insight into early responses to subinhibitory DOX in a 3D melanoma model, the fact that it uses only a single murine cell line limits its translational relevance. Further mechanistic validation and studies in more complex models mimicking the tumor microenvironment are needed, as the interplay of resident and infiltrated cells in TME is critical in shaping early adaptive responses to various chemotherapeutics ^75^.

This article contains supplemental data available at Repository: **Negrea et al, 2025;** doi: 10.17632/248c3t93fz.2 (^43^).

## Supporting information

https://opendata.openscience.ubbcluj.ro/drafts/248c3t93fz

## Disclosure: Funding information

This work was supported by L’Oréal - UNESCO “For Women in Science” [Fellowship Programme no. 914, 2020]; the UEFISCDI grant [PN-III-P2-2_1-PED-2021-0411, No. 659PED, 2022] granted to dr. Alina Sesarman.

## Declaration of generative AI in scientific writing

The authors acknowledge the use of generative AI (ChatGPT 3.5) in enhancing the readability and language of the manuscript. After using the tool, the authors thoroughly reviewed and edited the content, taking full responsibility for the final manuscript.

## Conflict of Interest

The authors have no conflict of interest.

## Abbreviations list

ABC: ATP-binding cassette transporters
Ackr3: atypical chemokine receptor 3
Angptl2: angiopoietin-like 2
APH: acid phosphatase assay
Apq1: Aquaporin 1
BAD: BCL-2-associated death promoter
Bard1: BRCA1-associated RING domain 1
BAX: BCL-2-associated X protein
BCL-2: B-cell lymphoma 2
BCL-W: BCL2-like 2
BIM: BCL-2-like 11
Bok: BCL2-related ovarian killer
bFGF: basic fibroblast growth factor
BID: BH3-interacting domain death agonist
Birc5: baculoviral IAP repeat-containing 5
Bst2: bone-marrow stromal cell antigen 2
Brip1: BRCA1-interacting protein C-terminal helicase 1
Bub1: BUB1 mitotic checkpoint serine/threonine kinase
BRAF: v-Raf murine sarcoma viral oncogene homolog B
Cdh6: cadherin-6
Ccn6: cellular communication network factor 6
cIAP-2: cellular inhibitor of apoptosis protein 2
Cdc45: cell-division cycle 45
CDC20: Cell division cycle 20
CDK1: cyclin-dependent kinase 1
CDKN1A/p21: cyclin-dependent kinase inhibitor 1A
p27: cyclin-dependent kinase inhibitor 1B
CD40/TNFRSF5: Cluster of Differentiation 40
CD40L: CD40 ligand
CTR1: control 1
CPM: counts per million
ECM: commercial extracellular matrix
Caspase-3: cysteine-aspartic acid protease-3
Caspase-8: cysteine-aspartic acid protease-8
CTLA-4: cytotoxic T-lymphocyte-associated protein 4
Cyclin G: Cyclin G1
cytoC: cytochrome C
DR6: Death Receptor 6
TOP2A: DNA topoisomerase II α
DOX: doxorubicin
DMEM: Dulbecco’s Modified Eagle Medium
DR6/TNFRSF21: Death Receptor 6
eotaxin-1: eotaxin-1
ECM: extracellular matrix
EGF: epidermal growth factor
EMT: epithelial– mesenchymal transition
Epha2: ephrin type-A receptor 2
EI24: etoposide-induced gene 2.4
Exo1: exonuclease 1
Fancd2: Fanconi anaemia complementation group D2
Fanci: Fanconi anaemia complementation group I
Fas: Fas cell-surface death receptor
Fas/apo-1: Fas cell-surface death receptor
FasL: Fas ligand
FCS: Fetal Calf Serum
FDR: false discovery rate
FR: fold-regulation value
FPKM: fragments per kilobase of transcript per million mapped reads
Foxm1: forkhead box M1
GSEA: Gene Set Enrichment Analysis
G2/M checkpoint: G2/M-phase cell-cycle checkpoint
G-CSF: granulocyte-colony-stimulating factor
GM-CSF: granulocyte–macrophage colony-stimulating factor
HTRA2: high-temperature-requirement protein A2
HSP27: heat-shock protein 27
HSP60: heat-shock protein 60
HSP70: heat-shock protein 70
HISAT2: Hierarchical Indexing for Spliced Alignment of Transcripts 2
Histone 1X: Histone 1X
IFN: interferon
IGF: insulin-like growth factor
IGF-1: insulin-like growth factor 1
IGF-2: insulin-like growth factor 2
IGFBP-1/2/3/4/5/6: insulin-like growth factor-binding protein 1,2,3,4,5,6
IL-16: interleukin-16
IKKβ: inhibitor of NF-κB kinase subunit β
Itga3: integrin α3
KEGG: Kyoto Encyclopedia of Genes and Genomes pathway
M-CSF: macrophage-colony-stimulating factor
MEK: MAPK/ERK kinase
MMP-2: matrix metalloproteinase-2
MMP-9: matrix metalloproteinase-9
Mad2l1: Mitotic arrest deficient 2-like 1
MCP-1: monocyte chemoattractant protein-1
mTORC1: mechanistic target of rapamycin complex 1
Nectin4: nectin-4
Nek2: NIMA-related kinase 2
NOD-like receptor: nucleotide-binding oligomerization domain-like receptor
NF-kB: Nuclear factor kappa B
Notch1: Notch homolog 1
Pidd1: p53-induced death-domain protein 1
Pen/Strep: penicillin/streptomycin
Plxdc2: plexin domain-containing protein 2
Parpbp: poly(ADP-ribose)-binding protein
p53: tumor protein p53
Pcdhβ5/7/8: protocadherin-β5,7,8
Polq: DNA polymerase θ
Psrc1: proline/serine-rich coiled-coil 1
Rad51: RAD51 recombinase
Rmi2: RecQ-mediated genome-instability protein 2
RPLP0: Ribosomal Protein Lateral Stalk Subunit P0
SMAC: second mitochondria-derived activator of caspase
Sesn: sestrin 2
Sfn: stratifin
Survivin: baculoviral inhibitor of apoptosis repeat-containing 5
Svep1: Sushi–von Willebrand factor, EGF and pentraxin domain-containing protein 1
TIMP-1/2: tissue inhibitor of metalloproteinase 1,2
Ticrr: TopBP1-interacting checkpoint and replication regulator
TMM: trimmed mean of M-values method
TRAIL R2: TRAIL receptor 2
Trex1: three-prime repair exonuclease 1
Tnfrsf19: tumor necrosis factor receptor superfamily member 19
TNF-α: tumour necrosis factor subunit α
TNF RI: tumour necrosis factor receptor I
TNFRSF1A/10B: tumour necrosis factor receptor superfamily member 1A, 10B
TPM: transcripts per million
TPO: thrombopoietin
VEGF: vascular endothelial growth factor
XIAP: X-linked inhibitor of apoptosis protein

## Notes

### Competing Interest Statement

The authors have declared no competing interest.

### Summary of Updates

We revised the preprint to improve clarity, structure, visual presentation and certain discussion points. The new version is more concise, mainly to improve overall readability. In addition, we improved several figures for more effective communication of the data.

## References

1. Huang, F., Santinon, F., Flores González, R. E. & Del Rincón, S. V. Melanoma Plasticity: Promoter of Metastasis and Resistance to Therapy. Front. Oncol. 11, 756001 (2021).

2. Fenton, S. E., Sosman, J. A. & Chandra, S. Resistance mechanisms in melanoma to immuneoncologic therapy with checkpoint inhibitors. Cancer Drug Resist. (2019) doi:10.20517/cdr.2019.28.

3. Huang, A. C. & Zappasodi, R. A decade of checkpoint blockade immunotherapy in melanoma: understanding the molecular basis for immune sensitivity and resistance. Nat. Immunol. 23, 660–670 (2022).

4. Lim, S. Y. et al. The molecular and functional landscape of resistance to immune checkpoint blockade in melanoma. Nat. Commun. 14, (2023).

5. Shi, H. et al. Acquired Resistance and Clonal Evolution in Melanoma during BRAF Inhibitor Therapy. Cancer Discov. 4, 80–93 (2014).

6. Ellerhorst, J. A. et al. Phase II trial of doxil for patients with metastatic melanoma refractory to frontline therapy. Oncol. Rep. (1999) doi:10.3892/or.6.5.1097.

7. Ugurel, Ms., et al. Clinical Phase II Study of Pegylated Liposomal Doxorubicin as Second-Line Treatment in Disseminated Melanoma. Oncol. Res. Treat. 27, 540–544 (2004).

8. Elliott, A. M. & Al-Hajj, M. A. ABCB8 Mediates Doxorubicin Resistance in Melanoma Cells by Protecting the Mitochondrial Genome. Mol. Cancer Res. 7, 79–87 (2009).

9. Licarete, E. et al. Overcoming Intrinsic Doxorubicin Resistance in Melanoma by Anti-Angiogenic and Anti-Metastatic Effects of Liposomal Prednisolone Phosphate on Tumor Microenvironment. Int. J. Mol. Sci. 21, 2968 (2020).

10. Panneerselvam, M., Bredehorst, R. & Vogel, C. W. Resistance of human melanoma cells against the cytotoxic and complement-enhancing activities of doxorubicin. Cancer Res. 47, 4601–7 (1987).

11. Vorobiof, D. A., Rapoport, B. L., Mahomed, R. & Karime, M. Phase II study of pegylated liposomal doxorubicin in patients with metastatic malignant melanoma failing standard chemotherapy treatment. Melanoma Res. 13, 201–3 (2003).

12. Licarete, E. et al. The prednisolone phosphate-induced suppression of the angiogenic function of tumor-associated macrophages enhances the antitumor effects of doxorubicin on B16.F10 murine melanoma cells in-vitro. Oncol. Rep. (2019) doi:10.3892/or.2019.7346.

13. Negrea, G. et al. Active Tumor-Targeting Nano-formulations Containing Simvastatin and Doxorubicin Inhibit Melanoma Growth and Angiogenesis. Front. Pharmacol. 13, (2022).

14. Galluzzi, L., Buqué, A., Kepp, O., Zitvogel, L. & Kroemer, G. Immunogenic cell death in cancer and infectious disease. Nat. Rev. Immunol. 17, 97–111 (2017).

15. Frank, N. Y. et al. ABCB5-Mediated Doxorubicin Transport and Chemoresistance in Human Malignant Melanoma. Cancer Res. 65, 4320–4333 (2005).

16. Hata, A. N. et al. Tumor cells can follow distinct evolutionary paths to become resistant to epidermal growth factor receptor inhibition. Nat. Med. 22, 262–269 (2016).

17. Pletz, N. et al. Doxorubicin-induced activation of NF-κB in melanoma cells is abrogated by inhibition of IKKβ, but not by a novel IKKα inhibitor. Exp. Dermatol. 21, 301–304 (2012).

18. Tian, Y. et al. PI3K/AKT signaling activates HIF1α to modulate the biological effects of invasive breast cancer with microcalcification. Npj Breast Cancer 9, 93 (2023).

19. Sharma, S. V. et al. A Chromatin-Mediated Reversible Drug-Tolerant State in Cancer Cell Subpopulations. Cell 141, 69–80 (2010).

20. Chauvistré, H. et al. Persister state-directed transitioning and vulnerability in melanoma. Nat. Commun. 13, 3055 (2022).

21. Müller, J. et al. Low MITF/AXL ratio predicts early resistance to multiple targeted drugs in melanoma. Nat. Commun. 5, (2014).

22. Karki, P., Angardi, V., Mier, J. C. & Orman, M. A. A Transient Metabolic State in Melanoma Persister Cells Mediated by Chemotherapeutic Treatments. Front. Mol. Biosci. 8, (2022).

23. Berthenet, K. et al. Failed Apoptosis Enhances Melanoma Cancer Cell Aggressiveness. Cell Rep. 31, 107731 (2020).

24. De Freitas, J. T. et al. Notch1 blockade by a novel, selective anti-Notch1 neutralizing antibody improves immunotherapy efficacy in melanoma by promoting an inflamed TME. J. Exp. Clin. Cancer Res. 43, (2024).

25. Sun, C. et al. Reversible and adaptive resistance to BRAF(V600E) inhibition in melanoma. Nature 508, 118–122 (2014).

26. Thakur, V. et al. Co-inhibition of Notch1 and ChK1 triggers genomic instability and melanoma cell death increasing the lifespan of mice bearing melanoma brain metastasis. J. Exp. Clin. Cancer Res. 44, (2025).

27. Yuan, Z. et al. Blockade of inhibitors of apoptosis (IAPs) in combination with tumor-targeted delivery of tumor necrosis factor-α leads to synergistic antitumor activity. Cancer Gene Ther. 20, 46–56 (2013).

28. Al-Hity, G. et al. An integrated framework for quantifying immune-tumour interactions in a 3D co-culture model. Commun. Biol. 4, 781 (2021).

29. Gonzalez-Fernandez, T., Tenorio, A. J., Saiz Jr, A. M. & Leach, J. K. Engineered Cell-Secreted Extracellular Matrix Modulates Cell Spheroid Mechanosensing and Amplifies Their Response to Inductive Cues for the Formation of Mineralized Tissues. Adv. Healthc. Mater. 11, (2022).

30. McDermott, M. et al. In vitro Development of Chemotherapy and Targeted Therapy Drug-Resistant Cancer Cell Lines: A Practical Guide with Case Studies. Front. Oncol. 4, (2014).

31. Srinivasan, B. & Lloyd, M. D. Dose–Response Curves and the Determination of IC _50_ and EC _50_ Values. J. Med. Chem. 67, 17931–17934 (2024).

32. Kuczynski, E. A., Sargent, D. J., Grothey, A. & Kerbel, R. S. Drug rechallenge and treatment beyond progression—implications for drug resistance. Nat. Rev. Clin. Oncol. 10, 571–587 (2013).

33. Lamichhane, A. & Tavana, H. Three-Dimensional Tumor Models to Study Cancer Stemness-Mediated Drug Resistance. Cell. Mol. Bioeng. 17, 107–119 (2024).

34. Shahi Thakuri, P., Luker, G. D. & Tavana, H. Cyclical Treatment of Colorectal Tumor Spheroids Induces Resistance to MEK Inhibitors. Transl. Oncol. 12, 404–416 (2019).

35. Friedrich, J. et al. A Reliable Tool to Determine Cell Viability in Complex 3-D Culture: The Acid Phosphatase Assay. SLAS Discov. 12, 925–937 (2007).

36. Longati, P. et al. 3D pancreatic carcinoma spheroids induce a matrix-rich, chemoresistant phenotype offering a better model for drug testing. BMC Cancer 13, 95 (2013).

37. Wang, Q., Lu, F. & Lan, R. RNA-sequencing dissects the transcriptome of polyploid cancer cells that are resistant to combined treatments of cisplatin with paclitaxel and docetaxel. Mol. Biosyst. 13, 2125–2134 (2017).

38. Pertea, M. et al. StringTie enables improved reconstruction of a transcriptome from RNA-seq reads. Nat. Biotechnol. 33, 290–295 (2015).

39. Benjamini, Y., Drai, D., Elmer, G., Kafkafi, N. & Golani, I. Controlling the false discovery rate in behavior genetics research. Behav. Brain Res. 125, 279–284 (2001).

40. Robinson, M. D., McCarthy, D. J. & Smyth, G. K. <tt>edgeR</tt> : a Bioconductor package for differential expression analysis of digital gene expression data. Bioinformatics 26, 139–140 (2010).

41. Kolberg, L. et al. g:Profiler—interoperable web service for functional enrichment analysis and gene identifier mapping (2023 update). Nucleic Acids Res. 51, W207–W212 (2023).

42. Subramanian, A. et al. Gene set enrichment analysis: A knowledge-based approach for interpreting genome-wide expression profiles. Proc. Natl. Acad. Sci. 102, 15545–15550 (2005).

43. Negrea, G.-G. et al. “Exploring the mechanisms of early acquired resistance to doxorubicin in melanoma spheroids”, Babeș-Bolyai University, V2, doi: 10.17632/248c3t93fz.2. (2025).

44. Rauca, V.-F. et al. Combination therapy of simvastatin and 5, 6-dimethylxanthenone-4-acetic acid synergistically suppresses the aggressiveness of B16.F10 melanoma cells. PLOS ONE 13, e0202827 (2018).

45. Leo, R. et al. Protein Expression Profiling Identifies Key Proteins and Pathways Involved in Growth Inhibitory Effects Exerted by Guggulsterone in Human Colorectal Cancer Cells. Cancers 11, 1478 (2019).

46. Sesarman, A. et al. Anti-angiogenic and anti-inflammatory effects of long-circulating liposomes co-encapsulating curcumin and doxorubicin on C26 murine colon cancer cells. Pharmacol. Rep. 70, 331–339 (2018).

47. Mrówczyńska, E., Mazurkiewicz, E. & Mazur, A. J. Gelatin Zymography to Detect Gelatinase Activity in Melanoma Cells. J. Vis. Exp. (2022) doi:10.3791/63278.

48. Fey, S. J., Korzeniowska, B. & Wrzesinski, K. Response to and recovery from treatment in human liver-mimetic clinostat spheroids: a model for assessing repeated-dose drug toxicity. Toxicol. Res. 9, 379–389 (2020).

49. Antonello, P. et al. ACKR3 promotes CXCL12/CXCR4-mediated cell-to-cell-induced lymphoma migration through LTB4 production. Front. Immunol. 13, (2023).

50. Enzler, T. et al. Cell-Selective Inhibition of NF-κB Signaling Improves Therapeutic Index in a Melanoma Chemotherapy Model. Cancer Discov. 1, 496–507 (2011).

51. Amiri, K. I. & Richmond, A. Role of nuclear factor-κ B in melanoma. Cancer Metastasis Rev. 24, 301–313 (2005).

52. Henriet, P. & Emonard, H. Matrix metalloproteinase-2: Not (just) a “hero” of the past. Biochimie 166, 223–232 (2019).

53. Holohan, C., Van Schaeybroeck, S., Longley, D. B. & Johnston, P. G. Cancer drug resistance: an evolving paradigm. Nat. Rev. Cancer 13, 714–726 (2013).

54. Filipiak-Duliban, A., Brodaczewska, K., Kajdasz, A. & Kieda, C. Spheroid Culture Differentially Affects Cancer Cell Sensitivity to Drugs in Melanoma and RCC Models. Int. J. Mol. Sci. 23, (2022).

55. Albertson, T. M. et al. DNA polymerase ε and δ proofreading suppress discrete mutator and cancer phenotypes in mice. Proc. Natl. Acad. Sci. 106, 17101–17104 (2009).

56. Keijzers, G. et al. Human Exonuclease 1 (EXO1) Regulatory Functions in DNA Replication with Putative Roles in Cancer. Int. J. Mol. Sci. 20, 74 (2018).

57. Karthikeyan, M. C. et al. Doxorubicin downregulates cell cycle regulatory hub genes in breast cancer cells. Med. Oncol. 41, 220 (2024).

58. Vera, J. et al. Chk1 and Wee1 control genotoxic-stress induced G2–M arrest in melanoma cells. Cell. Signal. 27, 951–960 (2015).

59. Liddiard, K., Aston-Evans, A. N., Cleal, K., Hendrickson, E. A. & Baird, D. M. POLQ suppresses genome instability and alterations in DNA repeat tract lengths. NAR Cancer 4, zcac020 (2022).

60. Singvogel, K. & Schittek, B. Dormancy of cutaneous melanoma. Cancer Cell Int. 24, 88 (2024).

61. Xiao, D., Dong, S., Yang, S. & Liu, Z. CKS2 and RMI2 are two prognostic biomarkers of lung adenocarcinoma. PeerJ 8, (2020).

62. Christowitz, C. et al. Mechanisms of doxorubicin-induced drug resistance and drug resistant tumour growth in a murine breast tumour model. BMC Cancer 19, (2019).

63. Rambow, F., Marine, J.-C. & Goding, C. R. Melanoma plasticity and phenotypic diversity: therapeutic barriers and opportunities. Genes Dev. 33, 1295–1318 (2019).

64. Ryl, T. et al. Cell-Cycle Position of Single MYC-Driven Cancer Cells Dictates Their Susceptibility to a Chemotherapeutic Drug. Cell Syst. 5, 237–250.e8 (2017).

65. Melnikova, V. O., Bolshakov, S. V., Walker, C. & Ananthaswamy, H. N. Genomic alterations in spontaneous and carcinogen-induced murine melanoma cell lines. Oncogene 23, 2347–2356 (2004).

66. Micheau, O. & Tschopp, J. Induction of TNF Receptor I-Mediated Apoptosis via Two Sequential Signaling Complexes of These Receptors Delivers a Powerful and Rapid Proapo-Ptotic Signal through a DD-Mediated Recruitment of the Adaptor Protein FADD and the Formation of the so-Called Complex I Fails to Induce the Expression of Antiapoptotic. Cell vol. 114 181–190 (2003).

67. Beaumont, K. A. et al. Cell Cycle Phase-Specific Drug Resistance as an Escape Mechanism of Melanoma Cells. J. Invest. Dermatol. 136, 1479–1489 (2016).

68. Gordon, E., Ravicz, J., Liu, S., Chawla, S. & Hall, F. Cell cycle checkpoint control: The cyclin G1/Mdm2/p53 axis emerges as a strategic target for broad-spectrum cancer gene therapy - A review of molecular mechanisms for oncologists. Mol. Clin. Oncol. (2018) doi:10.3892/mco.2018.1657.

69. Wu, S. & Singh, R. K. Resistance to Chemotherapy and Molecularly Targeted Therapies: Rationale for Combination Therapy in Malignant Melanoma. Curr. Mol. Med. 11, 553–563 (2011).

70. Deng, C. et al. Correction: TNFRSF19 Inhibits TGFβ Signaling through Interaction with TGFβ Receptor Type 1 to Promote Tumorigenesis. Cancer Res. 83, 1158–1158 (2023).

71. Endo, M. et al. Tumor Cell–Derived Angiopoietin-like Protein ANGPTL2 Is a Critical Driver of Metastasis. Cancer Res. 72, 1784–1794 (2012).

72. Meguenani, M. et al. Junctional adhesion molecule B interferes with angiogenic VEGF/VEGFR2 signaling. FASEB J. 29, 3411–3425 (2015).

73. Kunz, P. et al. Elevated ratio of MMP2/MMP9 activity is associated with poor response to chemotherapy in osteosarcoma. BMC Cancer 16, 223 (2016).

74. Spallarossa, P. et al. Matrix metalloproteinase-2 and -9 are induced differently by doxorubicin in H9c2 cells: The role of MAP kinases and NAD(P)H oxidase. Cardiovasc. Res. 69, 736–745 (2006).

75. Sun, Y. Tumor microenvironment and cancer therapy resistance. Cancer Lett. 380, 205–215 (2016).

