## Supplementary material for "Exploring the mechanisms of early acquired resistance to doxorubicin in melanoma spheroids": https://opendata.openscience.ubbcluj.ro/drafts/248c3t93fz

### Materials and methods

#### Quantitative Real-Time PCR (RT-qPCR)

RT-qPCR was conducted to confirm RNA-seq findings. 1 µg of total RNA was reverse transcribed into cDNA using iScript™ Reverse Transcription Supermix (#1708841, Bio-Rad Laboratories, Gladesville, Australia) in a 20 µL reaction volume, following manufacturer's instructions [1]. The reaction mix was incubated at 25°C for 5 minutes for priming, followed by reverse transcription at 46°C for 20 minutes, and enzyme inactivation at 95°C for 1 minute. Real-time quantitative PCR was then performed with SsoFast™ EvaGreen® Supermix (#172-5200, Bio-Rad Laboratories, Gladesville, Australia). Each 1 µL cDNA sample was amplified with 500 nM of specific primers. The primers sequences (purchased from Eurogentec, Seraing, Belgium) are available in **Supplementary Table 1** [2] (purchased from Eurogentec, Seraing, Belgium). The following setup was used for the amplification: enzyme activation at 95°C for 30 seconds, 40 cycles of denaturation at 95°C for 5 seconds, annealing at 55°C for 5 seconds, followed by a melting curve from 65°C to 95°C in 0.5°C increments. All reactions were run on the CFX96™ Real-Time System (C1000 Touch Thermal Cycler, Bio-Rad Laboratories, Gladesville, Australia) as per the manufacturer's protocol. Relative mRNA levels were calculated using the ΔΔCt method, with β-actin mRNA and ribosomal protein lateral stalk subunit P0 (Rplp0) serving as the internal references [3].

### Results

### Supplementary Figures

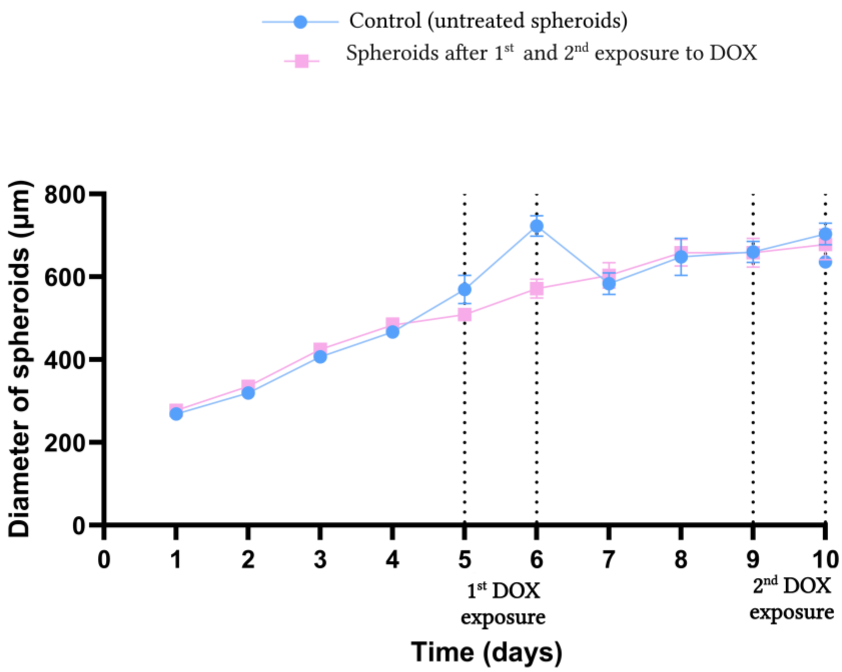


**Supplementary Figure 1. Graphic representation of spheroids diameter during experimental timeline (from day 1 to day 10).** B16.F10 melanoma spheroids exposed once (1st exposure), respectively, twice (2nd exposure) to DOX (IC30, 0.282 µM) compared to control, untreated spheroids. Results are presented as mean of three independent experiments (n=9).

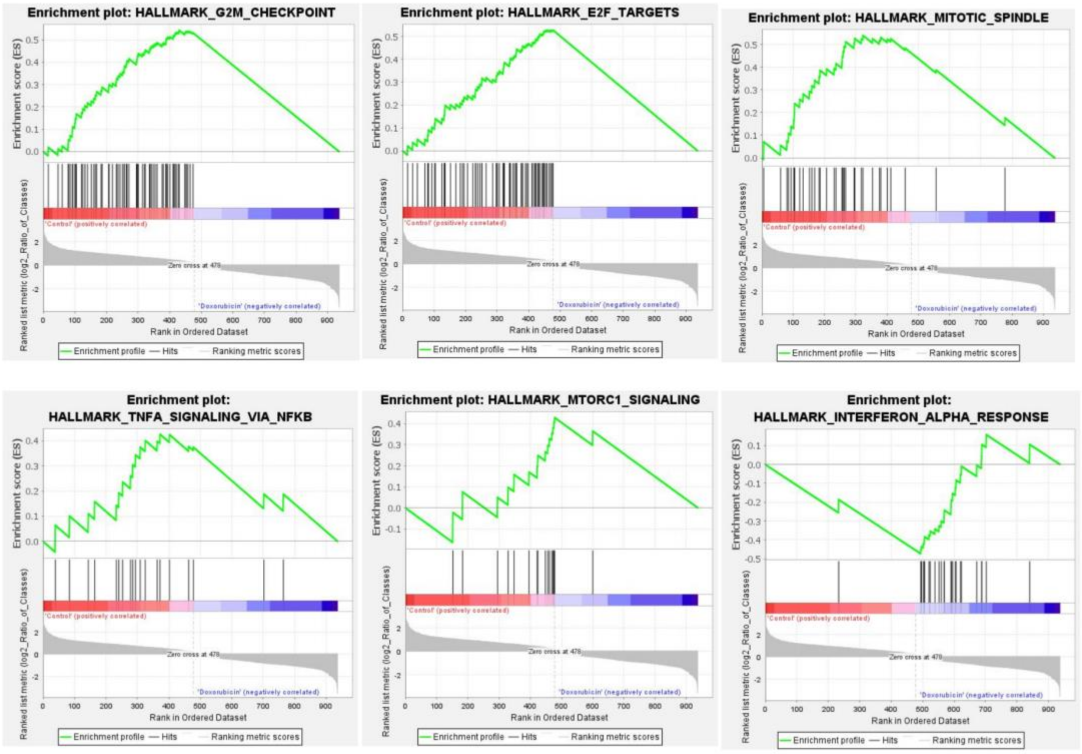


**Supplementary Figure 2. GSEA Enrichment plots of the corresponding six data sets enriched in GSEA.** Hallmark analysis showing the profile of the running ES Score and positions of gene set members on the rank-ordered list for significant gene sets enrichment at FDR < 20% and a nominal p-value < 5% in control (untreated)spheroids compared to spheroids after first exposure to DOX.

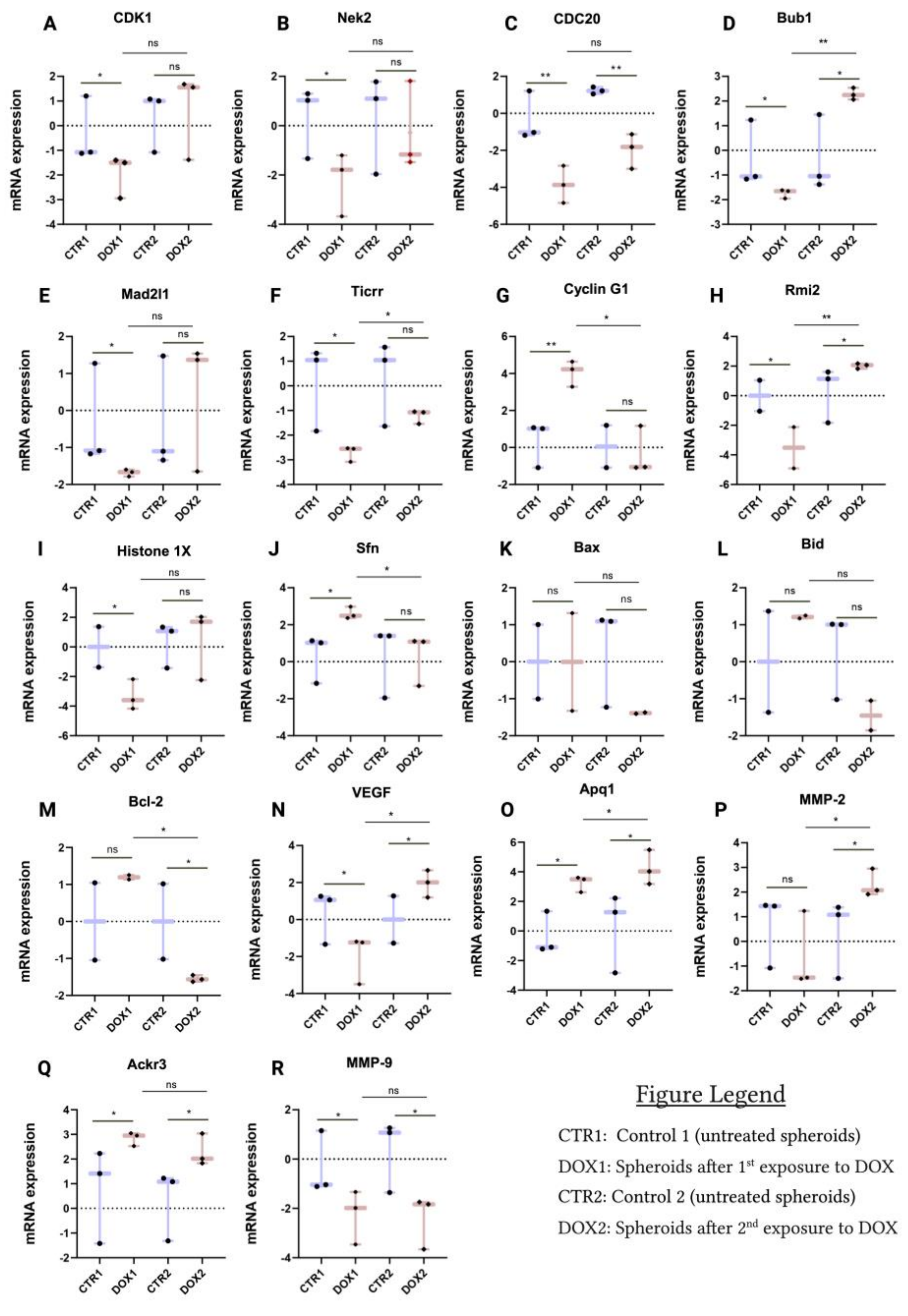


**Supplementary Figure 3. mRNA expression analysis in B16.F10 melanoma in spheroids after repeated exposure to DOX by RT-qPCR.** The figure presents the relative expression of 18 reference transcripts involved in the processes analyzed: **A-G**: cell cycle; **H-I**: DNA damage and repair; J-M: apoptosis; **N,O**: angiogenesis; **P-R**: migration in spheroids exposed to DOX (after 1st and 2nd exposures) compared to controls. β-actin and RPLP0 served as housekeeping genes in the qPCR analysis. The data are expressed as mean ± SD of at least two biological replicates; ns: not significant; **p* < 0.05, ***p* < 0.01.

**Supplementary tables** for **the** manuscript of **Negrea et. al, 2025** **"Exploring mechanisms of early acquired resistance to doxorubicin in melanoma spheroids"** are available **online** in **Mendeley Repository** as:

Negrea, Giorgiana-Gabriela; Pavel, Ilie Ovidiu; Balacescu, Loredana; Dume, Bogdan-Razvan; Licarete, Emilia; Rauca, Valentin-Florian; Patras, Laura; Meszaros, Szilvia; Dragan, Stefan; Toma, Vlad Alexandru; Banciu, Manuela; Sesarman, Alina (2025), "Exploring the mechanisms of early acquired resistance to doxorubicin in melanoma spheroids", Babeş-Bolyai University, V2, doi: 10.17632/248c3t93fz.2

**These data represent:**

1) Supplementary Table 1. RT-qPCR Primers sequences.
2) Supplementary Table 2. All DEGs.
3) Supplementary Table 3.1 GO and 3.2. KEGG analysis results.
4) Supplementary Table 4. GSEA analysis.
5) Supplementary Table 5. DEG in selected processes.

### References


1. Pflug, K. M., Lee, D. W., McFadden, K., Herrera, L. & Sitcheran, R. Transcriptional induction of NF-κB-inducing kinase by E2F4/5 facilitates collective invasion of GBM cells. *Scientific Reports* **13**, 13093 (2023).
2. Negrea, G.-G. *et al.* "Exploring the mechanisms of early acquired resistance to doxorubicin in melanoma spheroids", Babeş-Bolyai University, V2, doi: 10.17632/248c3t93fz.2. *Mendeley data, V2, doi: 10.17632/248c3t93fz.2* (2025).
3. Livak, K. J. & Schmittgen, T. D. Analysis of Relative Gene Expression Data Using Real-Time Quantitative PCR and the 2−ΔΔCT Method. *Methods* **25**, 402–408 (2001).
